# Beta Cells Deficient for *Renalase* Counteract Autoimmunity by Shaping Natural Killer Cell Activity

**DOI:** 10.1101/2024.02.29.582816

**Authors:** Kevin Bode, Siying Wei, Isabella Gruber, Stephan Kissler, Peng Yi

## Abstract

Type 1 diabetes (T1D) arises from autoimmune-mediated destruction of insulin-producing pancreatic beta cells. Recent advancements in the technology of generating pancreatic beta cells from human pluripotent stem cells (SC-beta cells) have facilitated the exploration of cell replacement therapies for treating T1D. However, the persistent threat of autoimmunity poses a significant challenge to the survival of transplanted SC-beta cells. Genetic engineering is a promising approach to enhance immune resistance of beta cells as we previously showed by inactivating of the *Renalase* (*Rnls*) gene. Here we demonstrate that *Rnls* loss-of-function in beta cells shape autoimmunity by mediating a regulatory Natural Killer (NK) cell phenotype important for the induction of tolerogenic antigen presenting cells. *Rnls*-deficient beta cells mediate cell-cell-contact-independent induction of hallmark anti-inflammatory cytokine Tgfβ1 in NK cells. In addition, surface expression of key regulatory NK immune checkpoints CD47 and Ceacam1 are markedly elevated on beta cells deficient for *Rnls*. Enhanced glucose metabolism in *Rnls* mutant beta cells is responsible for upregulation of CD47 surface expression. These findings are crucial to a better understand how genetically engineered beta cells shape autoimmunity giving valuable insights for future therapeutic advancements to treat and cure T1D.

**Graphical summary:** 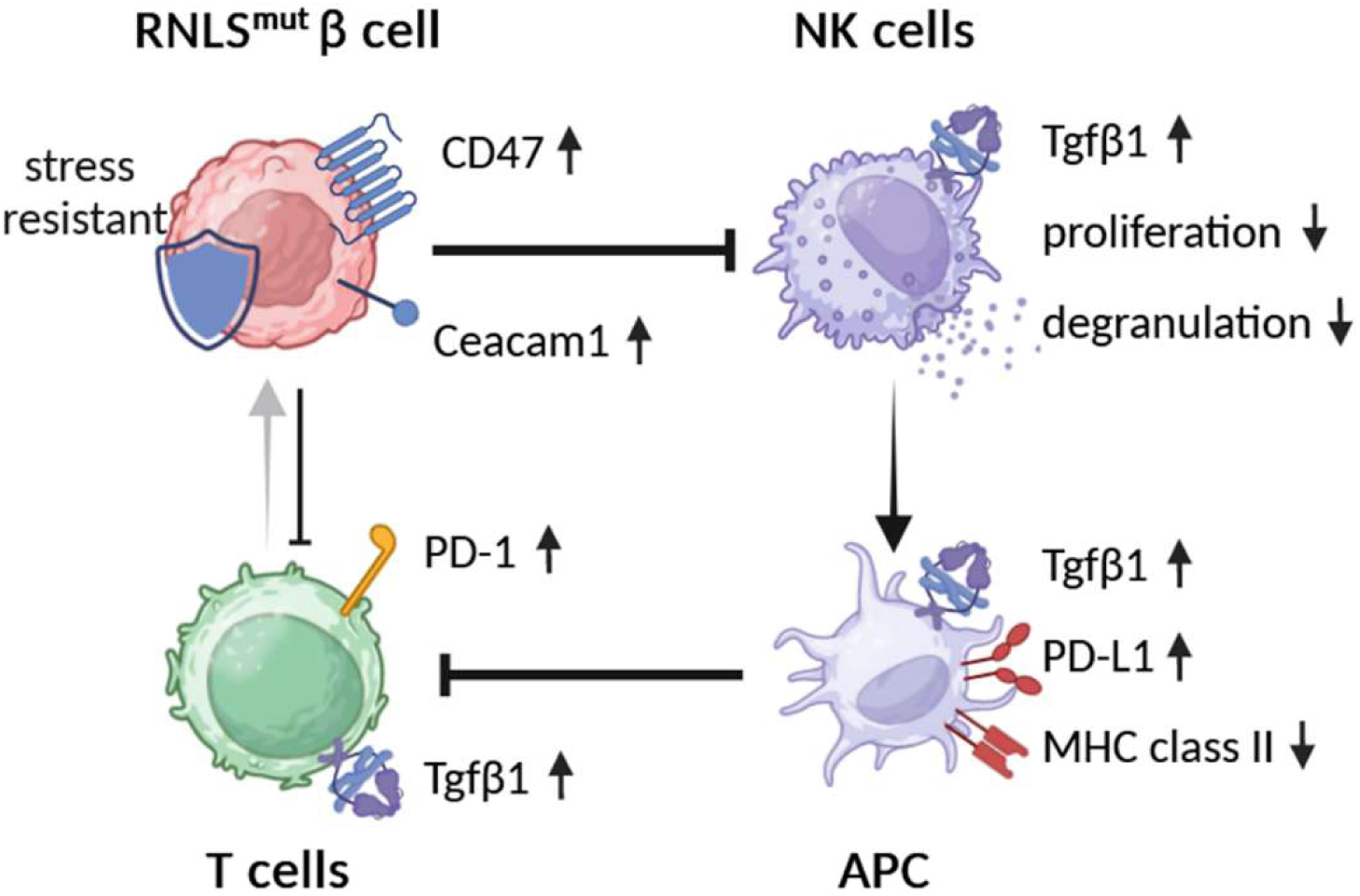

## INTRODUCTION

In Type 1 Diabetes (T1D), the autoimmune process leads to the selective destruction of insulin-producing beta cells within the pancreas. Upon extensive elimination of the majority of beta cell mass by autoimmunity, a curative strategy for T1D requires the replenishment of functional beta cell mass to completely restore the patient’s capacity for insulin production ^1^. The latest progress in the manufacturing of beta cells derived from stem cells (SC) has rendered beta cell replacement therapy a viable prospect ^2^. Despite these advancements, a significant hurdle in translating this strategy to clinical application lies in our current inability to protect beta cells against recurrent autoimmune attacks without resorting to broad immunosuppressive treatments. Exacerbating this challenge is the existing limitation in generating patient-specific SC-derived beta cells, underscoring the need to address both autoimmunity and alloimmune responses in any cellular therapy for T1D in the foreseeable future ^3^. To address these challenges, various research groups have started to genetically engineer SC-derived beta cells for enhanced resilience against immune-mediated destruction. Notably, most efforts focus on the deletion of genes encoding major histocompatibility complex (MHC) molecules mandatory for antigen presentation and activation of T cells, coupled with the incorporation of immune inhibitory ligands such as CD47 to prevent stimulation of innate immune cells. ^4–8^

In our pursuit of discovering novel immune-regulatory targets to protect beta cells from autoimmune destruction, we previously performed an unbiased genome-wide *in vivo* CRISPR screen, and found that Renalase (*Rnls*) deletion is able to protect beta cells from stress-induced cell death and autoimmunity ^9^. Beta cells lacking *Rnls* not only exhibit enhanced resilience to stress but also undergo comprehensive metabolic alterations favoring glycolysis ^9,10^. Moreover, we have shown previously that *Rnls*-deficient beta cells orchestrate localized shifts in the overall immune cell composition within the graft, primarily characterized by enhanced infiltration of CD4^+^ T cells, reduced numbers of natural killer (NK) cells, and by the accumulation of tolerogenic antigen-presenting cells (APC) defined by diminished MHC class II expression coupled with elevated levels of Programmed Cell Death 1 Ligand 1 (PD-L1). We also demonstrated that the survival advantage of beta cells lacking *Rnls* is attributed to the induced expression of PD-L1 on APC. Beta cells deficient for *Rnls* promote a significant transcriptional change in CD45^+^ immune cells within the graft towards hallmark anti-inflammatory genes such as Transforming Growth Factor beta 1 (Tgfβ1). ^10^ However, the immune cell population responsible for initiating the immuno-regulatory cross-talk to reduce autoimmunity against *Rnls*-deficient beta cell grafts remained elusive.

As Tgfβ1 is a known inducer of PD-L1 upregulation on APC in certain tumors as well as in pancreatic islet transplanation ^11^, and NK cells harbor as significant source of Tgfβ1 production ^12^, we have now investigated if graft-infiltrating NK cells play an important role for mediating the immuno-protective regulation of *Rnls* mutant beta cells. NK cells are not soley responsible for killing transformed or stressed target cells, but are also crucial to modulate the activation and phenotype of immune cells such as APC ^13^. Here, we show that depletion of NKp46^+^ innate lympoid cells in an experimental model of beta cell transplantation abrogated the induction of tolerogenic APC consequently preventing the survival advantage of beta cell grafts deficient for *Rnls*. Using *in vitro* co-culture systems we demonstrated that beta cells lacking *Rnls* are potent modulator of NK cell activation shown for both, mouse-derived NIT-1 beta cells and human SC-derived beta-like cells. Elevated cell surface expression of key NK cell inhibitory molecules on beta cells deficient for *Rnls* such as Cluster of differentiation 47 (CD47) and Ceacam1 are likley to play a major role in suppressing NK cell activation by cell-cell interaction. However, upregulation of Tgfβ1 in NK cells mediated by *Rnls* deletion in beta cells is independent of cell-cell contact demonstrating multifaced propsects of NK regulation mediated by *Rnls* deficieny in beta cells.

## RESULTS

### Protective immune-regulation by Rnls^mut^ beta cells requires NKp46^+^innate lymphoid cells

We previously investigated the consequences of *Rnls* deletion on immune cell infiltration and activation by transplantation of syngeneic mouse beta cell line NIT-1 into Non-obese diabetic (NOD) mice, a model of T1D. Single cell RNA sequencing of graft-infiltrating CD45^+^ cells revealed that *Rnls* mutant (Rnls^mut^) beta cells broadly influence immune cell activation and metabolism towards anti-inflammatory oncostatin M and Tgfβ1 expression, accompanied by enriched expression of genes important for glycolysis. In addition to elevated numbers of CD4^+^ T cells and tolerogenic PD-L1^+^ APC, beta cell grafts deficient for *Rnls* showed markedly reduced frequency of NK cells. Because we previously demonstrated that PD-L1 blockade abrogates the advantage of Rnls^mut^ beta cells to survive autoimmunity, and NK cells are known to modulate APC maturation and activation, we now investigated the role of NK cells in protection of Rnls^mut^ beta cells in more detail^10^. The gene expression profile of both, graft-infiltrating NK cells and closely related type 1 innate lymphoid cells (ILC1), demonstrated markedly increased expression of genes involved in pathways of cellular activation and inflammation (interferon responses, allograft rejection, etc.) when derived from WT beta cell grafts compared to Rnls^mut^ (Fig 1A and Supplemental Data Figure 1A-C), whereas ILC1 show downregulated gene expression of inflammatory cytokines interferon gamma (Ifng) and tumor necrosis factor alpha (Tnf). NK cells also upregulate gene expression of immune-regulatory Tgfβ1 when infiltrated in Rnls^mut^ beta cell grafts (Supplemental Data Figure 1A and B). Interestingly, the frequency of NK cells in Rnls^mut^ beta cell grafts was not only reduced in late stages ^10^, but also at earlier time points preceding the detection of graft weight difference between control and Rnls^mut^ (Supplemental Data Figure 1D and E; see Supplemental Data Figure 2 for gating strategy). Of note, the ILC1 frequency was not changed between the grafts and only represent a vast minority of infiltrating immune cells (≤ 1%) compared to NK cells (up to 14%) indicating that ILC1 likely does not play a major role in mediating the protective immune-regulation caused by Rnls^mut^ beta cells ^10^. These observations strengthened our hypothesis that NK cells might be crucial for regulating autoimmunity within Rnls^mut^ grafts. To investigate the importance of NK cells in shaping autoimmunity *in vivo*, we depleted NKp46^+^ innate lymphoid cells from autoreactive splenocytes before adoptive transfer into NIT-1 beta cell graft bearing mice (Figure 1B). The isogenic WT and Rnls^mut^ beta cells were transplanted subcutaneously (*s.c.*) on opposite flanks of the same recipient mice following intravenous (*i.v.*) injection of diabetogenic splenocytes similar as described before (Supplemental Data Figure 1D and E) ^9,10^. Strikingly, NKp46-depleted autoreactive splenocytes lost the ability to protect Rnls^mut^ cells from autoimmunity in both immunodeficient recipients, NOD-*Prkdc^scid^* (NOD.scid) mice and NOD.scid gamma (NSG) mice completely deficient for functional T, B and NK cells (Figure 1C). Whereas the accumulation of CD4^+^ T cells is independent of innate lymphoid cells, the reduced frequency of mature MHCII^high+^ APC in Rnls^mut^ grafts completely dependents on the presence of graft-infiltrating NKp46^+^ immune cells (Figure 1D and E). In the absence of innate lymphoid cells within Rnls^mut^ beta cell grafts, MHCII^high+^ APC lost the ability to upregulate the immune checkpoint PD-L1, responsible to forfeit the survival advantage over WT beta cells as shown before (Figure 1G and H; Bode *et al*., 2023). In summary, these observations clearly demonstrated that graft-infiltrating NKp46^+^ innate lymphoid cells are crucial for the induction of a tolerogenic APC phenotype important to reduce autoimmune responses to Rnls^mut^ beta cells.

**Figure 1:**
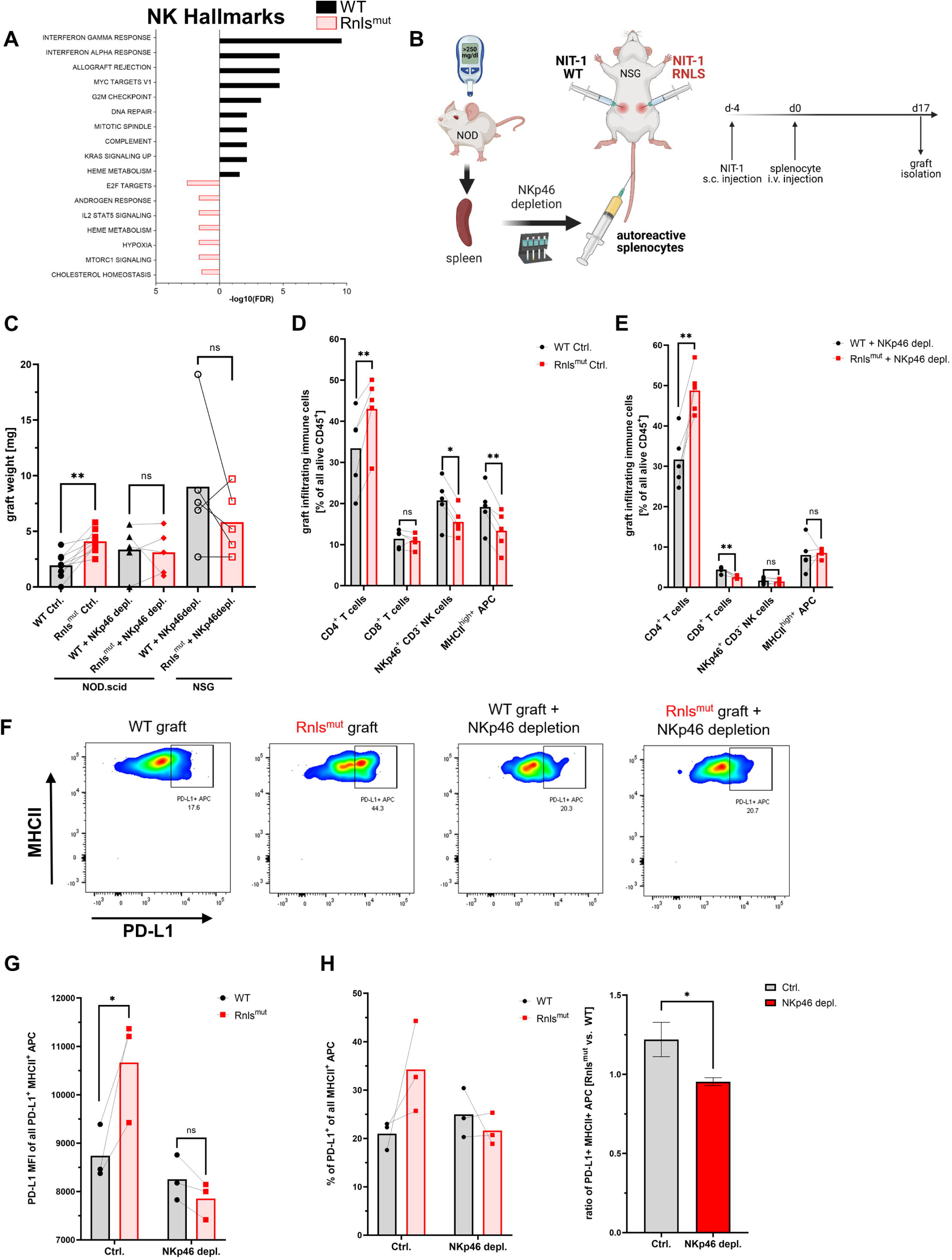
NKp46^+^ innate lymphoid cells are crucial for protective immune-regulation by *Renalase* mutant (Rnls^mut^) NIT-1 beta cells *in vivo*. (**A**) Hallmark gene expression analysis of natural killer (NK) cells derived from indicated NIT-1 beta cell grafts showing up to 10 most significantly changed pathways (-log10(FDR) ≥ 2) using 213 (Rnls^mut^) or 506 (WT) most significantly upregulated genes (p ≤ 0.05) as input. Raw data used in this figure were obtained from a 3’ gene expression single cell RNA sequencing experiment performed previously ^10^. (**B**) Schematic representation of the experimental design comparing paired WT and Rnls^mut^ NIT-1 beta cell grafts. Cells were injected *s.c.* into opposite flanks of indicated immunodeficient mice, followed by *i.v.* injection of autoreactive splenocytes 4 days later with or without preceding depletion of NKp46^+^ innate lymphoid cells. Grafts were harvested, scaled and immune cells were characterized by flow cytometry. (**c**) Weight of paired grafts from mice with or without depletion of NKp46**^+^** cells from autoreactive splenocytes are shown. Results represent the mean of nine (Ctrl. condition) or five (NKp46+ depletion) paired biological replicates from two combined independent experiments using NOD.scid or NSG recipient mice as indicated. (**D** and **E**) Quantification of immune cell subpopulations derived from five paired ctrl. NIT-1 beta cell grafts (**D**) or NKp46^+^ depleted grafts (**E**) as determined by flow cytometry. Results represent the mean of five paired biological replicates. (**F**-**H**) Representative flow cytometry data (**F**) or quantification (**G** and **H**) showing indicated expression level of PD-L1 on indicated immune cells derived from paired WT and *Rnls*-deficient grafts as determined by flow cytometry. Results represent the mean of five paired biological replicates. * p<0.05, ** p<0.01, ns p>0.05 (paired two-tailed t-test). Data obtained from scRNAseq (**A**) and from NKp46^+^ depletion experiments using NSG (**C-E**), or NOD.scid (**C**, **F**-**H**) recipient mice are derived from independent experiments. Data shown for the Ctrl. condition in (**C**) are partially derived from one previously performed experiment^10^ and from one newly performed independent experiment.

**Figure 2:**
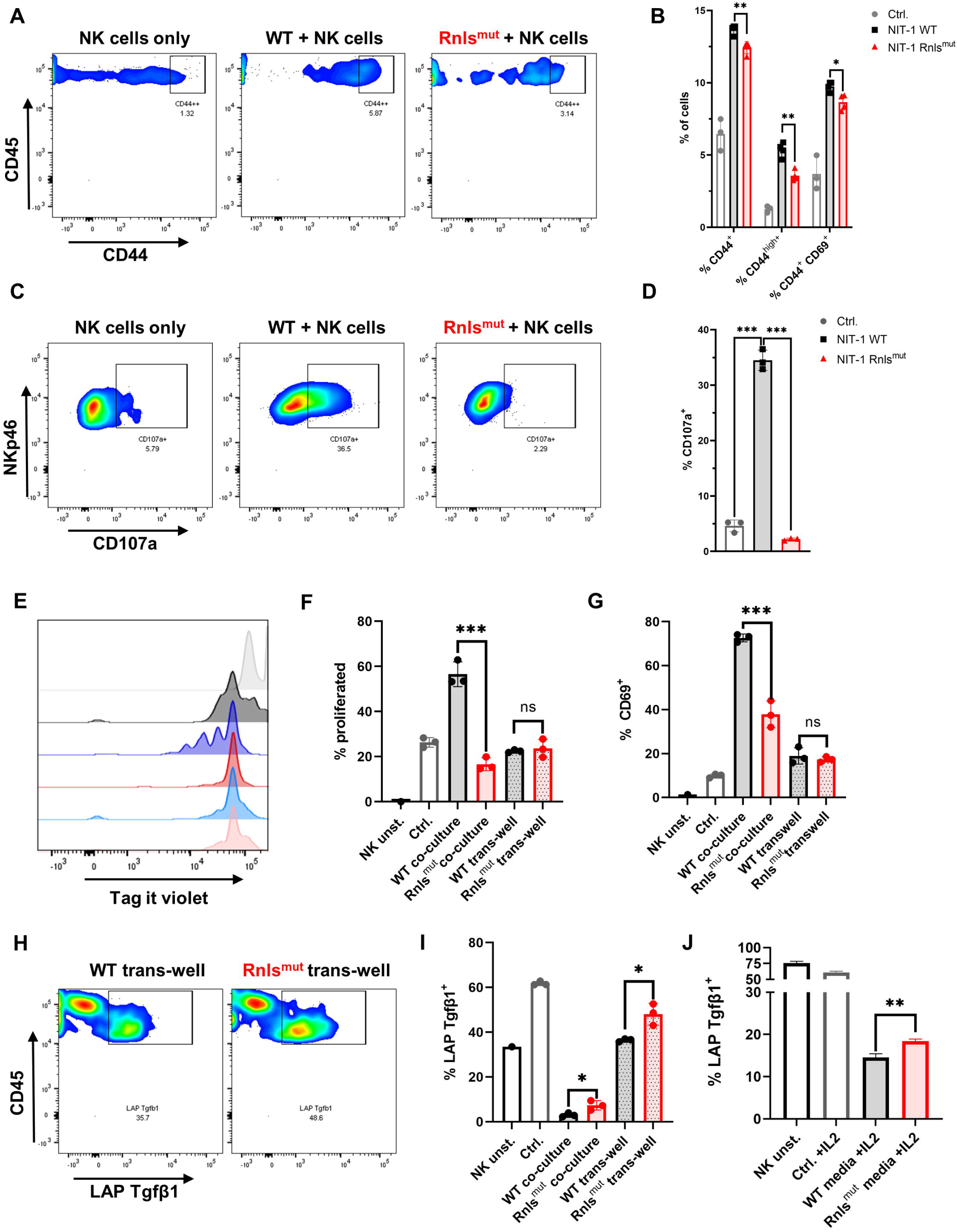
Rnls^mut^ NIT-1 beta cells broadly affect natural killer (NK) cell function *in vitro*. Interleukin 2 (IL2) stimulated primary mouse NK cells were co-cultured with indicated NIT-1 beta cells for 72h. (**A**-**D**) Data derived from single dose IL2 (20 ng/ml) treated co-culture experiments showing representative flow cytometry plots (**A** and **C**) or related quantifications (**B** and **D**) of indicated activation marker (**A** and **B**) or cytotoxic activity marker CD107a (**C** and **D**). (**E**-**I**) NK cells were repetitively stimulated with IL2 (100 ng/ml) every 24h together with indicated NIT-1 cells in co-culture or trans-well setting. (**E** and **F**) Representative flow cytometry data (**E**) or quantifications (**F**) showing NK cell proliferation of indicated conditions. (**G**) Quantification of surface expression of activation marker CD69 on NK cells. (**H** and **I**) Representative flow cytometry data derived from trans-well conditions (**H**) or quantifications of indicated conditions (**I**) showing expression level of membrane-bound latency-associated peptide (LAP)/Tgfβ1 expression on NK cells. (**J**) Conditioned media experiments showing LAP/Tgfβ1 surface expression on NK cells. Results represent the mean ± SD from one out of two (**A**, **B**, **E**-**I**) or three (**C**, **D** and **J**) independent experiments (n=3). *** p<0.001, ** p<0.01, * p<0.05, ns p>0.05, (unpaired, two-tailed t-test).

### Rnls^mut^ NIT-1 beta cells shape NK cell activation towards a regulatory phenotype

After showing that NKp46^+^ innate lymphoid cells are important to mediate protective immune-regulation leading to prolonged graft survival of Rnls^mut^ beta cells, we subsequently investigated if Rnls^mut^ beta cells directly regulate the activation of innate lymphoid cells. Hence, we purified splenic NKp46^+^ cells (further declared as NK cells as ILC1 only represents about 5-10% of all NKp46^+^ cells in the spleen)^14^ by negative selection to collect “untouched” cells for functional *in vitro* co-culture experiments. Activation of NK cells was achieved by supplementation of interleukin-2 (IL2), well-known to enhance proliferation and effector function of NK cells ^15^. Co-culture with Rnls^mut^ NIT-1 beta cells significantly reduce NK cell activation indicated by impaired expression of activation marker CD44 and CD69 compared to WT, but did not completely abolish NK cell activation (Figure 2A and B). However, the surface expression of degranulation marker CD107a (also known as LAMP1) on NK cells, a molecule that predicts cytotoxic activity ^16^, is strikingly abolished when co-cultured with Rnls^mut^ NIT-1 beta cells (Figure 2C and D). The ability of NK cells to proliferate following repeated IL2 stimulations was also completely prevented when co-cultured with Rnls^mut^ NIT-1 beta cells. This effect was solely dependent on cell-cell contact as separation of NK cells and Rnls^mut^ beta cells in trans-well abrogated the anti-proliferative function of Rnls^mut^ beta cells (Figure 2E and F). The cell-cell contact dependency on NK cell regulation by Rnls^mut^ NIT-1 beta cells also became evident for regulation of activation marker CD69 (Figure 2G). As described above, differential gene expression analysis of graft-infiltrated NK cells show enriched Tgfβ1 when derived from Rnls^mut^ NIT-1 beta cell grafts (Supplemental Data Figure 1A). To investigate if Tgfβ1 is also upregulated on protein level, we stained NK ells for latency-associated peptide (LAP) representing membrane-bound Tgfβ1. Indeed, NK cells co-cultured with Rnls^mut^ NIT-1 beta cells demonstrated elevated Tgfβ1 cell surface expression compared to WT NIT-1 beta cells. In contrast to the modulation of proliferation, Rnls^mut^ NIT-1 beta cells drive Tgfβ1 expression independent of cell-cell contact (Figure 2H-J). That Rnls^mut^ NIT-1 beta cells significantly upregulates Tgfβ1 expression in NK cells by soluble factors were demonstrated by two different methods, in a trans-well assay (Figure 2H and I) and by supplementation of conditioned media derived from Rnls^mut^ NIT-1 beta cells in comparison to supernatant of WT cells (Figure 2J). In summary, Rnls^mut^ NIT-1 beta cells directly shape NK cell activity in multiple modes, dependent and independent of cell-cell contact. The presence of NIT-1 beta cells deficient for *Rnls* causes NK cells to fail to proliferate as well as to lose cytotoxic activity in response to IL2, but at the same time elevate the expression level of anti-inflammatory Tgfβ1.

### NIT-1 beta cells deficient for Rnls show upregulation of inhibitory immune checkpoint molecules important to regulate NK cell activity

We demonstrated that Rnls^mut^ NIT-1 beta cells are potent modulator of NK cell activation (Figure 1 and 2). To explore how *Rnls*-deficient beta cells might influence the activity of NK cells, we re-analyzed bulk RNA sequencing comparing differential gene expression data comparing WT and Rnls^mut^ NIT-1 cells^10^. Rnls^mut^ NIT-1 beta cells show an enrichment of genes involved in immune-regulatory interactions between a lymphoid and a non-lymphoid cell (Supplemental Data Figure 1F) indicating that upregulated expression of inhibitory cell surface molecules may regulate cell contact-dependent NK cell stimulation. Next, we selected all candidates from top 1500 most significant upregulated genes in Rnls^mut^ NIT-1 cells that are known to be involved in modulation of immune cell activity according to the GSEA-MSigDB data base (https://www.gsea-msigdb.org). In addition to elevated expression of CD44 and Itga4 that is important for cell-cell adhesion process^17,18^, Rnls^mut^ NIT-1 beta cells also upregulate key immune checkpoint surface molecules such as Ceacam1 (also known as CD66a), CD47 (also known as integrin associated protein), and CD200 (also known as OX-2), which are all eminent potent inhibitors of NK cell activation (Figure 3A) ^19–22^. To validate if elevated mRNA expression correspond to higher protein expression, we performed cell surface staining for these three key inhibitory NK cell ligands analyzed by flow cytometry. Rnls^mut^ NIT-1 beta cells show significant upregulated surface expression of CD47 for both, percentage of cells and for mean fluorescence intensity (MFI, Figure 3B and C). The surface expression of Ceacam1 was also significantly upregulated in terms of Ceacam1^high+^ expressing cells, whereas Ceacam1 MFI show a tendency of elevated expression on Rnls^mut^ NIT-1 beta cells (Figure 3E and F). Although the percentage of cells positive for cell surface expression of CD200 was not dramatically changed between WT and Rnls^mut^, the MFI of CD200 on Rnls^mut^ NIT-1 beta cells was significantly upregulated (Figure 3H and I). These observations indicate that Rnls^mut^ NIT-1 beta cells could shape NK cell activation at least partially by upregulation of multiple inhibitory NK cell ligands on their cell surface.

**Figure 3:**
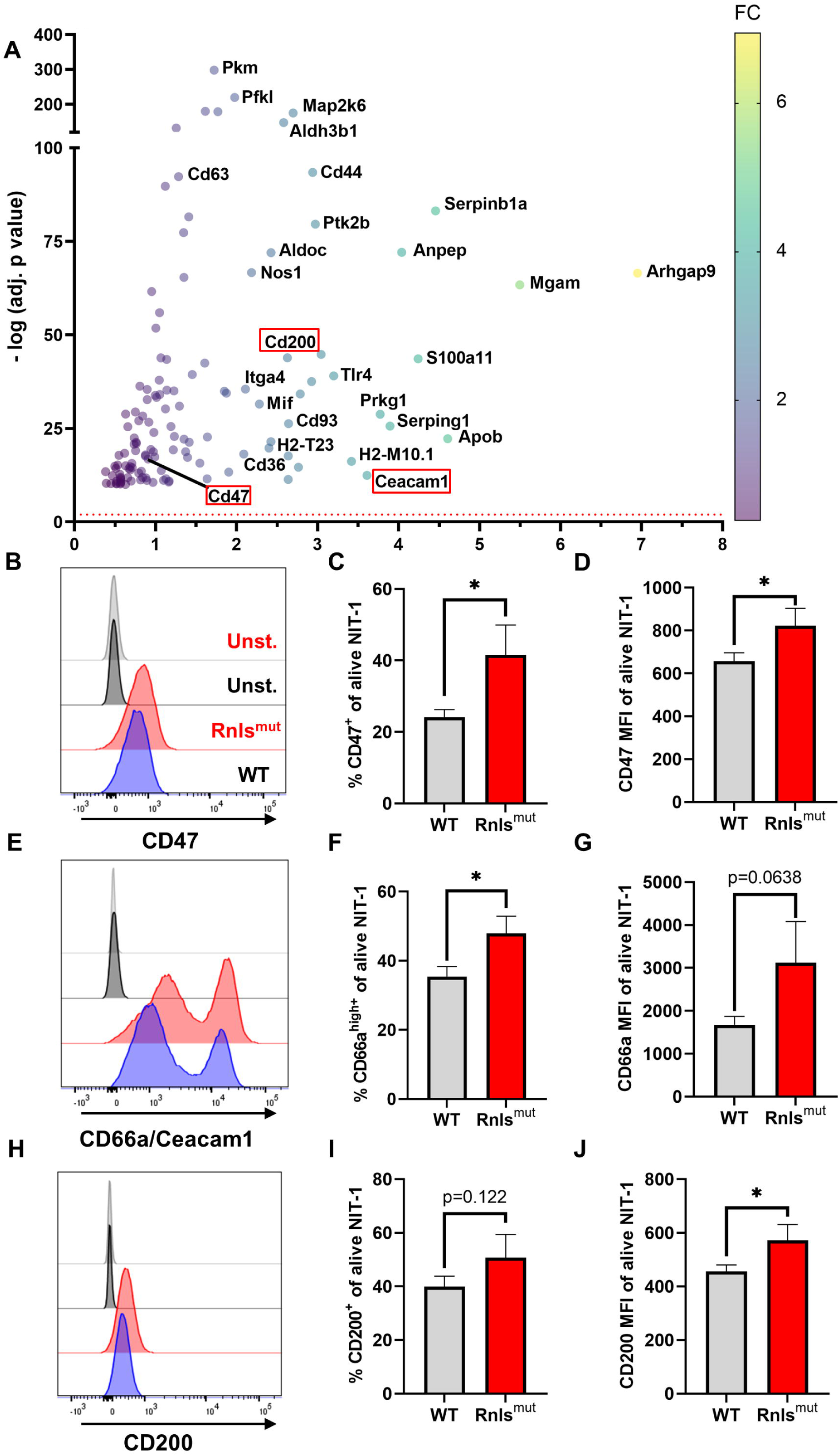
Rnls^mut^ NIT-1 beta cells show upregulated expression of key NK inhibitory ligands on their cell surface. (**A**) RNA sequencing data showing top 1500 upregulated immune cell function related genes in Rnls^mut^ NIT-1 beta cells in comparison to WT control. Cell surface molecules known to reduce NK cell activity are outlined with a red box. The red dotted line indicates the threshold for significantly changed genes. The fold change of gene expression is represented by colored dots from low (purple) to high (yellow). Raw data used in this figure were obtained from a bulk RNA sequencing experiment performed previously ^10^. (**B**-**J**) Representative histogram plots or quantifications of indicated inhibitory NK cell ligands expressed on the cell surface of Rnls^mut^ and WT NIT-1 cells characterized by flow cytometry. Results represent the mean ± SD from one out of three independent experiments (n=3). * p<0.05, (unpaired, two-tailed t-test).

In a first attempt to investigate why Rnls^mut^ cells upregulate the expression of NK inhibitory molecules we wondered if the changes in Rnls^mut^ beta cell metabolism towards increased glycolysis^10^ could influence the cell surface expression of CD47. As CD47 expression has been described to be similarly regulated like the immune checkpoint PD-L1 ^23^, and PD-L1 has been described to be positively regulated by glycolysis ^24,25^, we treated NIT-1 cells with 2-deoxy-d-glucose (2DG) to inhibit glucose metabolism. Strikingly, 2DG treatment dramatically downregulated the surface expression of CD47 on both, alive WT and Rnls^mut^ NIT-1 cells (Supplemental Data Figure 3A-D). Whereas treatment with low concentration of 2DG (1 mM) for 48 h decreased the percentage of CD47-expressing Rnls^mut^ cells to about the level of untreated WT cells, small amounts of 2DG have no effect on the percentage of CD47 expression on WT cells (Supplemental Data Figure 3A and C). This observation indicates that enhanced glycolysis in Rnls^mut^ beta cells is responsible for upregulation of CD47 surface expression.

### Human RNLS^mut^ SC-derived beta-like cells (SCBC) recapitulate key features of NK cell regulation

Rnls^mut^ NIT-1 beta cells strongly modulate NK cell activity towards a regulatory phenotype. To make sure that our observations are not restricted to mouse-derived NIT-1 beta cells, we also analyzed human iPSC-derived beta-like cells (SCBC) lacking the *RNLS* gene ^9^ for NK regulatory characteristics (Figure 4A). Intracellular staining for the beta cell marker C-peptide and NKX6.1 shows that about 72% of all cells successfully differentiated into beta-like cells independent of the genotype as described before (Figure 4B and C) ^9^. Rnls^mut^ NIT-1 beta cells show elevated surface expression of NK inhibitory molecules CD47, Ceacam1 and CD200 (Figure 3). In line with Rnls^mut^ NIT-1 beta cells, RNLS^mut^ SCBC also demonstrate significantly upregulated expression of NK inhibitory ligands CD47 and CD66a/c/e whereas, in contrast to NIT-1 beta cells, CD200 surface expression was not elevated on RNLS^mut^ SCBC (Figure 4D and E, Supplemental Data Figure 4A-F). We co-cultured WT and RNLS^mut^ with allogenic peripheral blood mononuclear cells (PBMC) stimulated with IL2 for 3 days (Figure 4A). Although surface expression of NK cell activation marker were only modestly changed in this highly stimulatory allogenic setting (data not shown), RNLS^mut^ SCBC significantly upregulates the immune-regulatory cytokine TGFβ1 on human CD3^-^ CD56^+^ NK cells (Figure 4F). These findings clearly show that human beta cells lacking *RNLS* acquire key characteristics to induce a regulatory NK cell phenotype. Because we previously demonstrated that Tgfβ1 expression is elevated on other immune cells such as CD3^+^ CD4^+^ T cells when infiltrating in Rnls^mut^ beta cell grafts ^10^, we stained human CD3^+^ T cells for membrane-bound TGFβ1 as well. Indeed, PBMC-derived CD3^+^ T cells show significantly elevated surface expression of LAP/TGFβ1 when co-cultured with RNLS^mut^ SCBC compared to WT SCBC (Figure 4G). Finally, we tested if RNLS^mut^ SCBC have higher resistance to immune-mediated destruction by co-culture with IL2 stimulated allogenic PBMC. The total number of alive CD45 negative SCBC cells was significantly higher for RNLS^mut^ SCBC compared to WT indicating that the protective immune-regulatory effects of *RNLS*-deficient beta cells also translate, at least to some extent, to the human system (Figure 4H). Our results highlight that knocking out a single gene in beta cells can drive multifaceted modulation on the immunogenicity of beta cells to prevent autoimmunity.

**Figure 4:**
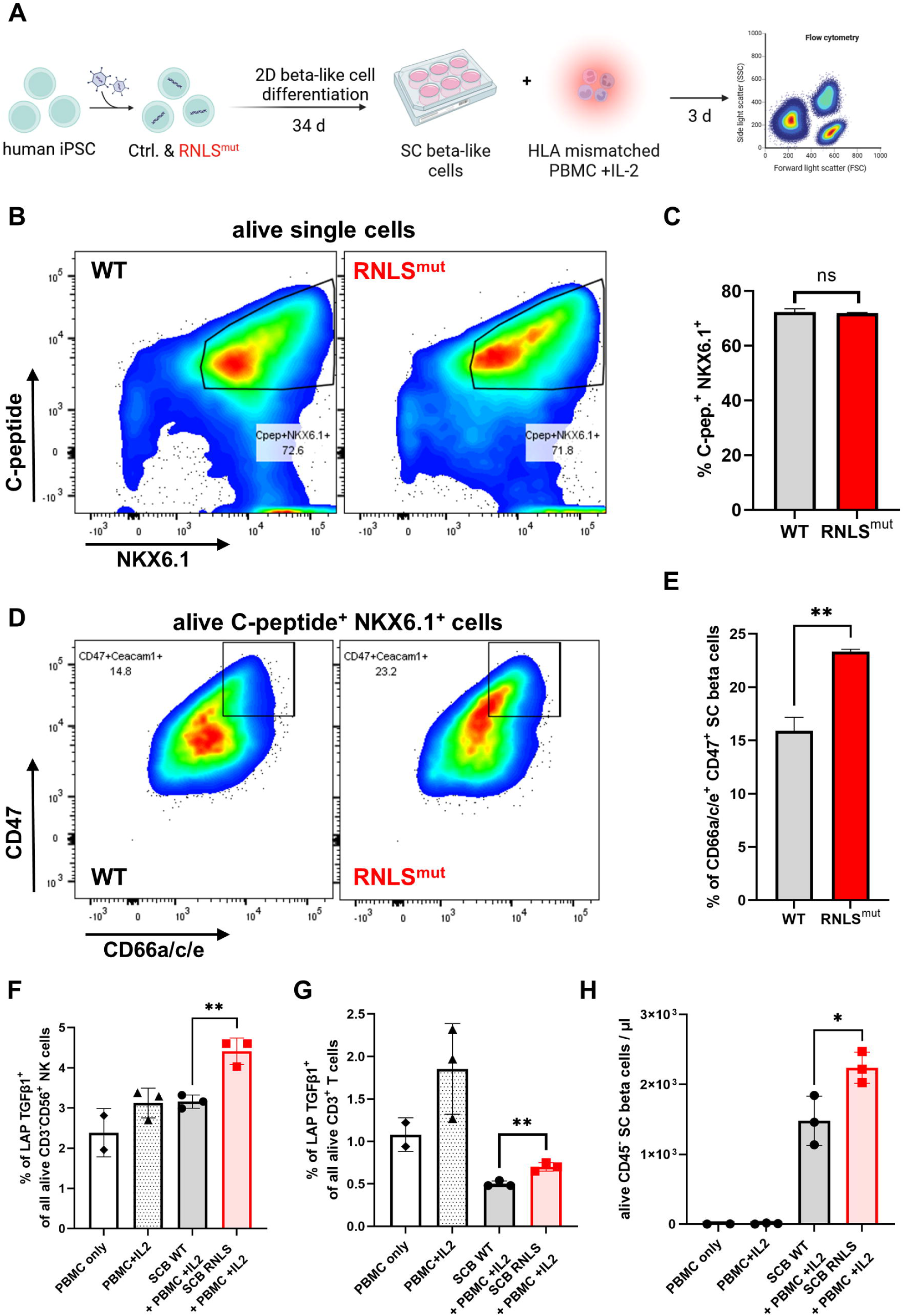
Rnls^mut^ stem cell (SC)-derived beta-like cells (SCBC) recapitulate key immuno-regulatory features to modulate NK cell activity. (**A**) Schematic representation of the experimental design for SCBC differentiation and for human leukocyte antigen (HLA) mismatched PBMC co-culture approach comparing WT and RNLS-deficient SCBC. (**B** and **C**) Representative flow cytometry plots or quantification of indicated beta cell marker of WT and RNLS^mut^ SCBC. (**D** and **E**) Representative flow cytometry plots or quantification of indicated NK inhibitory ligands on the cell surface of C-peptide^+^ NKX6.1^+^ WT and RNLS^mut^ SCBC. (**F** and **G**) Quantification of latency-associated peptide (LAP)/TGFβ1 surface expression on indicated immune cell types following 72h incubation with IL2-stimulated HLA mismatched PBMC in co-culture with WT or RNLS^mut^ SCBC. (**H**) Quantification of indicated alive CD45^-^ SCBC following IL2-stimulated HLA mismatched PBMC co-culture for 72h. Results represent the mean ± SD (n=2-3). * p<0.05, ** p<0.01, ns p>0.05 (unpaired two-tailed t-test).

## DISCUSSION

In this work we provide compelling evidence supporting the critical involvement of NK cells in orchestrating the immune-regulatory milieu surrounding Rnls^mut^ beta cells. In our previous study we already observed a marked reduction in NK cell frequency within Rnls^mut^ beta cell grafts compared to WT counterparts, suggesting a potential link between NK cells and the protective effects conferred by Rnls^mut^ beta cells ^10^. Here, further characterization of graft-infiltrating NK cells revealed distinct gene expression profiles, with NK cells derived from Rnls^mut^ beta cell grafts exhibiting upregulated expression of immune-regulatory Tgfβ1, reinforcing their role in mitigating autoimmune responses. Dysregulation of Tgfβ1 signaling has been implicated in various autoimmune disorders, highlighting its crucial role in maintaining immune homeostasis and preventing autoimmunity ^26^. Notably, depletion of NKp46^+^ cells abrogated the protective effects of Rnls^mut^ beta cells against autoimmunity, underscoring the indispensability of innate lymphoid cells, especially NK cells, in mediating immune regulation within the beta cell graft microenvironment.

Our previous work had established that PD-L1 blockade abrogates the survival advantage of Rnls^mut^ beta cells, implicating the PD-1/PD-L1 checkpoint as a crucial mediator of immune regulation in beta cell graft survival. In extension of our previous finding, our current observations elucidate the role of NKp46^+^ innate lymphoid cells in shaping the phenotype of APCs within the beta cell grafts. We observed a significant reduction in the frequency of mature MHCII^high+^ APCs within Rnls^mut^ beta cell grafts, which was completely dependent on the presence of graft-infiltrating NKp46^+^ immune cells. Enhanced secretion of Tgfβ1 derived from NK cells most likely induce the tolerogenic APC phenotype in Rnls^mut^ beta cell grafts. It is well established that Tgfβ1 inhibits dendritic cell maturation and upregulation of MHCII expression underscoring the intricate crosstalk between NK cells and APC we have observed here ^27^. In the context of lung cancer, the interplay between NK cells and APC has also been noted to have immune-regulatory implications, including impairments in MHCII expression and in modulation of immune checkpoint pathways ^28^. Of note, and in line with our observations, Tgfβ1 has been described as potent inducer of PD-L1 expression on APC in pancreatic islet transplantation ^11^. This suggests that NK cells play a pivotal role in shaping the tolerogenic phenotype of APC, thereby contributing to the overall immune-regulatory environment conducive for survival of beta cell grafts.

In this study, *in vitro* co-culture experiments provided mechanistic insights into the direct modulation of NK cell activity by Rnls^mut^ beta cells. Beta cells deficient for *Rnls* exhibited the ability to attenuate NK cell activation and cytotoxicity, while concurrently promoting the expression of anti-inflammatory Tgfβ1. Interestingly, Rnls^mut^ beta cells also upregulated inhibitory immune checkpoint molecules such as CD47, Ceacam1, and CD200, further corroborating their role in dampening NK cell activity and immune responses ^19–22,29^. Overexpression of CD47 on hypo-immunogenic MHCI/II-deficient islets have already been described to play a significant role in the inhibition of NK cells to prevent rejection of engrafted islets in humanized mice ^30^. However, other surface molecules or immune-modulatory alterations such as cellular metabolism in Rnls^mut^ beta cells may contribute to the inhibition of NK cell activity as well. The results from this study indicates that enhanced glucose metabolism in Rnls^mut^ beta cells contributes to elevated expression of CD47, as the inhibition of glycolysis by 2DG treatment strongly reduced cell surface expression of CD47. It has been shown that PD-L1 expression is regulated by glucose metabolism ^24^, but it has not been described before that CD47 surface expression is regulated similarly by glucose metabolism. It is known that the expression of PD-L1 and CD47 is often regulated by similar mechanisms such as by stimulation with IFNγ ^23^. However, future studies have to investigate the molecular mechanism how glucose metabolism is linked to the upregulation of CD47 surface expression on beta cells.

Moreover, our findings extend to human beta-like cells derived from induced pluripotent stem cells, highlighting the translational relevance of our observations. RNLS^mut^ SCBC exhibited similar patterns of NK cell regulation and upregulated expression of inhibitory immune checkpoint molecules CD47 and CD66a/c/e. In addition to mouse-derived NIT-1 cells, RNLS^mut^ SCBC also mediate the induction of TGFβ1 on both, human NK cells and T cells ^10^, further affirming the conservation of immune-modulatory mechanisms across species. This is in line with our previous study where we demonstrated that RNLS^mut^ SCBC is resistant to stress similar to *Rnls*-deficient NIT-1 cells ^9^.

Overall, our study elucidates the intricate interplay between Rnls^mut^ beta cells and NKp46^+^ NK cells in orchestrating protective immune regulation, unveiling potential targets for therapeutic intervention in autoimmune diabetes. By unraveling the mechanisms underlying immune modulation within the beta cell microenvironment, our findings pave the way for the development of novel strategies aimed at preserving beta cell function and improving graft survival in T1D.

## RESEARCH DESIGN AND METHODS

### Cell lines and stem cell-derived beta-like cell differentiation

NIT-1 cells were obtained from ATCC (cat # CRL-2055). Cells were maintained in DMEM, high glucose, pyruvate (cat # 11-995-073; Gibco), supplemented with 2 mM L-glutamine (cat # 25-030-081; Gibco),10% FCS (cat # 10-082-147; Gibco), 50 μM 2-Mercaptoethanol (cat # 60-24-2; Sigma-Aldrich), and penicillin/streptomycin (cat # 15140122; Thermo Fisher Scientific) in a 37 °C incubator with 5% CO_2_. Rnls^mut^ NIT-1 cells were generated previously (Bode *et al.*, 2023). In brief, *Rnls* gRNA (5′-CTACTCCTCTCGCTATGCTC-3′, MGLibA_46009) were cloned into lentiCRISPR v.2 vector. Lentivirus containing non-targeting or *Rnls* gRNA were used to establish the cell lines. The *Rnls* mutation was confirmed by deep-sequencing analysis (MGH DNA Core Facility) and quantitative PCR (qPCR; data not shown). Human induced pluripotent stem cells (iPSCs) obtained from Local donor 3 with type 1 diabetes were carefully maintained and differentiated following robust protocols tailored for beta cell differentiation^31^. Cultured in planar form using mTeSR medium (cat# 85850; Stem Cell Technologies), iPSCs were feed with daily media changes to sustain optimal growth conditions. Employing a 2-D differentiation protocol ^32^, iPSCs were seeded at stage 0 onto culture plates at a density of 0.8*10^5^ cells/cm^2^, supplemented with 10μM Y-27632 to enhance cell survival and adherence. Quality control measures were rigorously implemented at each differentiation stage, utilizing immunofluorescence (IF) or flow cytometry analyses with specific differentiation markers to assess differentiation efficiency and cellular identity. For effective cell dissociation, TrypLE Express (cat # 12604013; Gibco) treatment was administered at 37°C for 10 minutes to disrupt cell-cell adhesions, facilitating the generation of single cell suspensions. Following dissociation, cells were promptly fixed and stained as per established procedures^33^ to visualize cellular markers indicative of beta cell differentiation. Quality control at stage 6 involved the use of antibodies targeting C-Peptide and Nkx6.1 (see Table 1). Ethical approval for all human cell experiments was meticulously obtained from the Harvard University Institutional Review Board (IRB) and the Embryonic Stem Cell Research Oversight (ESCRO) committees, underscoring our commitment to ethical research practices and compliance with regulatory guidelines. RNLS^mut^ iPSCs from an individual with T1D (Local donor 3) were obtained and generated by the Melton Lab as described before^9^.

**Table 1.** Antibodies used for flow cytometry.

| **Antibody specificity** | **reactivity** | **Clone** | **Cat #** | **Vendor** |
| --- | --- | --- | --- | --- |
| C-Peptide Alexa Fluor 647 | human | U8-424 | 565831 | BD Biosciences |
| CD3 Brilliant Violet 785 | mouse | 17A2 | 100232 | Biolegend |
| CD3 Brilliant Violet 605 | mouse | 17A2 | 100237 | Biolegend |
| CD3 PE-Cy5 | human | HIT3a | 300310 | Biolegend |
| CD4 APC | mouse | RM4-5 | 100516 | Biolegend |
| CD8 FITC | mouse | 53-6.7 | 100706 | Biolegend |
| CD11b APC-Cy7 | mouse | M1/70 | 101226 | Biolegend |
| CD11c Brilliant Violet 711 | mouse | N418 | 117349 | Biolegend |
| CD19 Pacific Blue | mouse | 1D3/CD19 | 152416 | Biolegend |
| CD44 APC-Cy7 | mouse | IM7 | 103028 | Biolegend |
| CD44 PE | human | C44Mab-5 | 397504 | Biolegend |
| CD45 PE-Cy7 | mouse | 30-F11 | 103114 | Biolegend |
| CD45 PE-Cy7 | human | 2D1 | 368532 | Biolegend |
| CD47 APC | mouse | Miap301 | 127514 | Biolegend |
| CD47 PE-Cy7 | human | CC261 | 323113 | Biolegend |
| CD56 Brilliant Violet 605 | human | HCD56 | 318334 | Biolegend |
| CD66a | mouse | Mab-CC1 | 134540 | Biolegend |
| CD66a/c/e Brilliant Violet 421 | human | ASL-32 | 342313 | Biolegend |
| CD69 PE-Cy5 | mouse | H1.2F3 | 104510 | Biolegend |
| CD69 Brilliant Violet 421 | human | FN50 | 310930 | Biolegend |
| CD107a FITC | mouse | 1D4B | 121606 | Biolegend |
| CD200 (OX2) APC | mouse | OX90 | 123810 | Biolegend |
| CD200 (OX2) Brilliant Violet 711 | human | OX-104 | 329222 | Biolegend |
| F4/80 FITC | mouse | BM8 | 123108 | Biolegend |
| I-A<sup>k</sup> (MHC class II) PE | mouse | 10-3.6 | 109908 | Biolegend |
| LAP/TGF-β1 PE | mouse | TW7-16B4 | 141404 | Biolegend |
| LAP/TGF-β1 FITC | human | S20006A | 300010 | Biolegend |
| Ly-6C Brilliant Violet 421 | mouse | HK1.4 | 128032 | Biolegend |
| Ly-6G Brilliant Violet 785 | mouse | 1A8 | 127645 | Biolegend |
| NKp46 Brilliant Violet 711 | mouse | 9E2 | 331936 | Biolegend |
| NKp46 PerCP/Cyanine5.5 | mouse | 29A1.4 | 137610 | Biolegend |
| Nkx6.1 PE | human | R11-560 | 563023 | BD Biosciences |
| PD-L1 PerCP/Cyanine5.5 | mouse | 10F.9G2 | 124334 | Biolegend |

### Bulk RNA sequencing

Bulk RNAseq of WT and Rnls^mut^ NIT-1 cells were performed previously (Cai *et al.*, 2020). In brief, triplicates of 2 x 10^6^ NIT-1 cells were used for RNA isolation using Zymo Quick-RNA miniprep plus kit (cat # R1058; Zymo Research), following the manufacturers protocol. RNA libraries were prepared, and subsequently quality control tested by Novogene Corporation Inc. Sequencing was performed on an Illumina NovaSeq 6000 sequencing system for >20 million read data output (Novogen Corporation Inc.). For enrichment analysis of immune system-related genes (according to GSEA-MSigDB, https://www.gsea-msigdb.org) the 1500 most upregulated genes in *Rnls* mutant vs. WT control were used (Figure 3A).

### NK cell / NIT-1 beta cell co-culture experiments

NK cells were obtained from spleens of 8-16 wks old non diabetic NOD/ShiLtJ mice purchased from Jackson laboratory (strain #: 001976; 3-4 spleens were used per experiment to collect sufficient amount of NK cells) using the NKp46^+^ NK Isolation Kit (cat # 130-115-818; Miltenyi Biotec). For co-culture experiments, 10^4^ NIT-1 cells were seeded in each well of a 96-well flat bottom (trans-well assays) or round bottom (co-culture assays) plate in 100 μl of NK cell medium (DMEM, high glucose, pyruvate (cat # 11-995-073; Gibco), supplemented with 10% FCS (cat # 10-082-147; Gibco),2 mM L-glutamine (cat # 25-030-081; Gibco), and penicillin/streptomycin (cat # 15140122; Thermo Fisher Scientific)) following incubation for 3 d in a 37 °C incubator with 5% CO_2_. In trans-well assays NK cells were fluorescently labeled using *Tag-it Violet* proliferation dye according to the manufacturers’ protocol (cat # 425101; Biolegend) and 5*10^4^ NK cells were added to NIT-1 cell-containing 96-well plates either in direct co-culture or on a trans-well insert (cat # 3380; Corning). Recombinant mouse (m)IL-2 (cat # 575406; Biolegend) was added to stimulate NK cells with a total of 20ng/ml-100ng/ml final concentration as indicated. For trans-well assays 100 ng/ml mIL-2 was freshly added daily to achieve stronger NK cell proliferation. To investigate NK cell proliferation, NK cells were stained with Tag-it Viole Proliferation and Cell Tracking Dye (cat # 425101; Biolegend) according to manufacturer’s instructions. Conditioned media experiments were performed by the addition of medium derived from WT or Rnls^mut^ NIT-1 cells with ratio of 1:1 (conditioned medium : NK cell medium). Additional conditioned medium was supplemented following 24h (ratio 3:1) and 48h (4:1) each time in combination with repetitive IL-2 stimulation (100 ng/ml) for a total incubation time of 72h. In all experiments cells were co-cultured in a 37 °C incubator with 5% CO_2_ for 72 h, then collected and stained with indicated antibodies listed in Table 1. PI viability dye was used for dead cell staining and flow cytometry was performed on an LSR II instrument (BD Biosciences). Data were analyzed using *FlowJo* v.10.6.1 (FlowJo LLC).

### PBMC / SCBC co-culture experiments

Peripheral blood mononuclear cells (PBMC) collected from allogenic donors in EDTA tubes were isolated using Lymphoprep density gradient medium (cat # 07801; Stem Cell Technologies). A total number of 5*10^6^ PBMC were transferred into each well of a 6-well plate with fully confluent WT or RNLS^mut^ SCBC in 3ml of HEPES buffered PBMC medium ((RPMI-1640 with penicillin/streptomycin, L-glutamine and HEPES (cat # ABI-Custom American BioInnovations), supplemented with 10% human AB serum (cat # H5667; Millipore Sigma), and 100 ng/ml recombinant human (h)IL-2 (cat # 589104; Biolegend)) following incubation at 37 °C with 5% CO_2_. Following co-culture for 72 h, cells were detached with Trypsin-EDTA (cat # 25-200-056; Gibco) and filtered using a 70 μm pore size strainer two times to obtain a single cell solution. Collected cells were stained with indicated antibodies listed in Table 1. The Quantification of alive CD45^-^ SCBC was performed by using Precision Count Beads (cat # 424902; Biolegend). PI viability dye was used for dead cell staining and flow cytometry was performed on an LSR II instrument (BD Biosciences). Data were analyzed using *FlowJo* v.10.6.1 (FlowJo LLC).

### Splenocyte transfer mouse model and graft isolation

WT and *Rnls^mut^* NIT-1 beta cells (10^7^ each) were injected subcutaneously (*s.c.*) into opposite flanks of the same immunodeficient NOD.Cg-Prkdc^scid^/J (NOD; strain #: 001303) or NOD.Cg-Prkdc^scid^ Il2rg^tm1Wjl^/SzJ (NSG; starin #: 005557) mice purchased from Jackson laboratory. Four days later, autoreactive splenocytes from recently diabetic NOD mice were isolated and red blood cell were lysed using 5 ml of ACK lysing buffer (cat # A1049201; Thermo Fisher Scientific) for 4 min at RT. Lysis was stopped using 5 ml of PBS/ 10 % FCS followed by two washing steps using PBS only. Splenocytes were filtered two times in total through a strainer with 70 μm pore size to obtain a single cell solution. For NK cell depletion, NKp46-expressing splenocytes were removed by using the anti-NKp46 MicroBead kit (cat # 130-095-390; Miltenyi Biotec) according to manufacturers’ instructions. Splenocytes (10^7^/mouse) were injected intravenously (*i.v.*) into NIT-1 cell-bearing NOD.scid/NSG recipient mice to transfer autoimmune beta cell killing for a total of 17 days. NIT-1 beta cell graft isolation was performed as described before ^10^. In brief, Grafts were isolated and scaled on an analytical balance 17 days after splenocytes transfer. Grafts were cut into small pieces and digested using HEPES buffered RMPI 1640 (cat # R4130-10L; Sigma-Aldrich) supplemented with 1mg/ml Collagenase D ( cat # 11088858001; Sigma-Aldrich), 20 μg/ml DNase I (cat # EN0521; Thermo Fisher Scientific), 2% FCS and 50 μg/ml Lipopolysaccharide neutralizing agent Polymyxin B sulfate (cat # 1405-20-5; Sigma-Aldrich) for 45 min at 37°C while shaking by maximum speed on a heating block. Digested grafts were further disaggregated and filtered through a strainer with 70 μm pore size two times to obtain a single cell solution. For flow cytometry analysis cells were stained with the antibodies listed in Table 1 in two separate panels (panel 1: T cells and NK cells; panel 2: Myeloid cells and B cells).

### Single cell RNA sequencing (scRNAseq)

The scRNAseq experiment was performed in our previous study (Bode *et al.*, 2023). In brief, WT or *Rnls* mutant graft-infiltrating immune cells were FACS sorted (CD45^+^ PI^-^). Individual samples were labeled using hashtag antibodies and scRNAseq was performed on pooled samples using the Chromium Next GEM Single Cell 3’ GEM, Library & Gel Bead Kit v3.1 (cat # PN-1000213; 10 x Genomics) according to manufacturer’s instructions. Samples were super-loaded with 40,000 cells per reaction. The hashtag oligo library (HTO) was generated separately as described previously (https://citeseq.files.wordpress.com/2019/02/cell_hashing_protocol_190213.pdf). Illumina NovaSeq 6000 with approximately 1.1 billion reads was used to sequence the gene expression library, while the HTO library was sequenced separately by Illumina NextSeq with about 130 million reads total each. For further details on QC and analysis procedure please refer to our previous work (Bode *et al.*, 2023). Gene set enrichment analyses of graft-infiltrating NK cells and ILC1 were performed by using the molecular signatures of hallmark gene sets provided by the MSigDB database (http://www.gsea-msigdb.org/). The indicated number of differential expressed genes within the indicated immune cell population derived from WT or Rnls^mut^ NIT-1 beta cell grafts were used as input.

### Flow cytometry

All staining procedures were performed in PBS/10 % FCS. Single cell suspensions were pre-incubated with FcR-blocking antibody solution (cat # 156604; Biolegend) for 5 min, then stained with antibodies against indicated cell surface molecules (Table 1) for 20 min on ice diluted in FcR-blocking solution. Appropriate isotype controls were purchased from Biolegend. Dead cells were excluded using Propidium Iodide (PI, cat # R37169; Thermo Fisher Scientific) or fixable viability dye eFluor 450 (cat # 65-0863; eBiosciences). For intracellular staining the Transcription Factor Staining Buffer Set (cat # 00-5523-00; Thermo Fisher Scientific) was used according to manufacturer’s instruction. For the 2DG experiment to inhibit gluose metabolism in NIT-1 cells, 2*10^5^ cells of indicated genotypes were seeded into a 24-well plate for 48 h with or without addition of indicated concentrations of 2DG or sterile dH_2_O as control. Cell surface staining of indicated molecules were performed as described above. Data were acquired on an LSRII instrument (BD Biosciences) and analyzed with *FlowJo* software v.10.6.1 (FlowJo LLC). All data are shown using log-scale axes.

## Acknowledgments.

We thank Taylor Stewart (Joslin Diabetes Center) and Erica P. Cai (Lilly Diabetes Center of Excellence at the Indiana Biosciences Research Institute) for providing essential reagents. In some parts of the manuscript OpenAI (ChatGPT) was just used to optimize texting.

## Author Contributions

K.B. performed transplantation, flow cytometry, SCBC PBMC experiments, SCBC QC, analysed data, wrote the manuscript and supervised this work. S.W. generated mutant iPSC, differentiated SCBC and performed SCBC QC. I.G. purified NK cells, performed functional co-culture experiments, flow cytometry, and analysed data. S.K. and P.Y. provided supervision for some parts of this work and wrote parts of the manuscript. All authors reviewed the final manuscript and approved submission. K.B. is the guarantor of this work and, as such, had full access to all the data in the study and takes responsibility for the integrity of the data and the accuracy of the data analysis.

## Data availability

Datasets from our scRNAseq experiments are available from the Gene Expression Omnibus (https://www.ncbi.nlm.nih.gov/geo/) under accession number GSE226361 as described before ^10^.

