## Supplemental Data for "Beta Cells Deficient for *Renalase* Counteract Autoimmunity by Shaping Natural Killer Cell Activity"

#### Supplemental Data Figure Legends

**Supplemental Data Figure 1: Rnls$^{mut}$ graft-infiltrating NK cells and ILC1 show altered frequency and gene expression profiles.** (**A** and **B**) Volcano plot showing differential gene expression (DGE) between NK cells (**A**) and type 1 innate lymphoid cells (ILC1; **B**) derived from *Rnls*-deficient and WT NIT-1 beta cell grafts. Raw data used in this figure were obtained from a 3' gene expression single cell RNA sequencing experiment performed previously [10]. Altered expression of key cytokines are marked by red dots. The gray line indicates the threshold for significant changed gene expression (-log10 (p value) ≥ 1.3). (**C**) Hallmark gene expression analysis showing the 10 most significantly changed pathways (-log10 (FDR) ≥ 2) in ILC1. For ILC1 derived from Rnls$^{mut}$ and WT grafts, 430 or 355 significantly upregulated genes were used as input, respectively. (**D** and **E**) WT and Rnls$^{mut}$ NIT-1 beta cell cells were injected *s.c.* into opposite flanks of immunodeficient NOD.scid mice, followed by *i.v.* injection of autoreactive splenocytes 4 days later. Grafts were harvested before any noticeable differences in graft weights were detected (**E**) and immune cells were characterized by flow cytometry (**D**; See Supplemental Figure 2 for representative flow cytometry plots showing gating strategy). Results represent the mean of six paired biological replicates. *** $p<0.001$, ** $p<0.01$, ns $p>0.05$ (paired two-tailed t-test). (**F**) RNA sequencing data showing enrichment plot for immune-regulatory interactions between a lymphoid and a non-lymphoid cell of Rnls$^{mut}$ NIT-1 beta cells in comparison to WT control. Raw data used in this figure were obtained from a bulk RNA sequencing experiment performed previously [10].

**Supplemental Data Figure 2: Gating strategy to identify indicated cell subsets of graft-infiltrating immune cells.** Representative flow cytometry plots showing the overall gating strategy are depicted.

**Supplemental Data Figure 3: CD47 surface expression on NIT-1 beta cells is regulated by glucose metabolism.** Quantifications of CD47 expression on the cell surface of WT (**A** and **B**) and $Rnls^{mut}$ (**C** and **D**) NIT-1 beta cells following treatment with indicated concentrations of 2-deoxy-d-glucose (2DG) characterized by flow cytometry. Results represent the mean ± SD from one out of three independent experiments (n=3). **** $p<0.001$, *** $p<0.001$, ** $p<0.01$, * $p<0.05$, ns $p>0.05$, (unpaired, two-tailed t-test).

**Supplemental Data Figure 4: SCBC deficient for *Rnls* show elevated CD47 and CD66a/c/e surface expression.** Representative histogram plots or quantifications of indicated inhibitory NK cell ligands expressed on the cell surface of $RNLS^{mut}$ and WT induced pluripotent stem cell (iPSC)-derived beta-like cells (SCBC) characterized by flow cytometry. Results represent the mean ± SD (n=2-3). ** $p<0.01$, * $p<0.05$, ns $p>0.05$ (unpaired, two-tailed t-test).

### Supplemental Data Figure 1

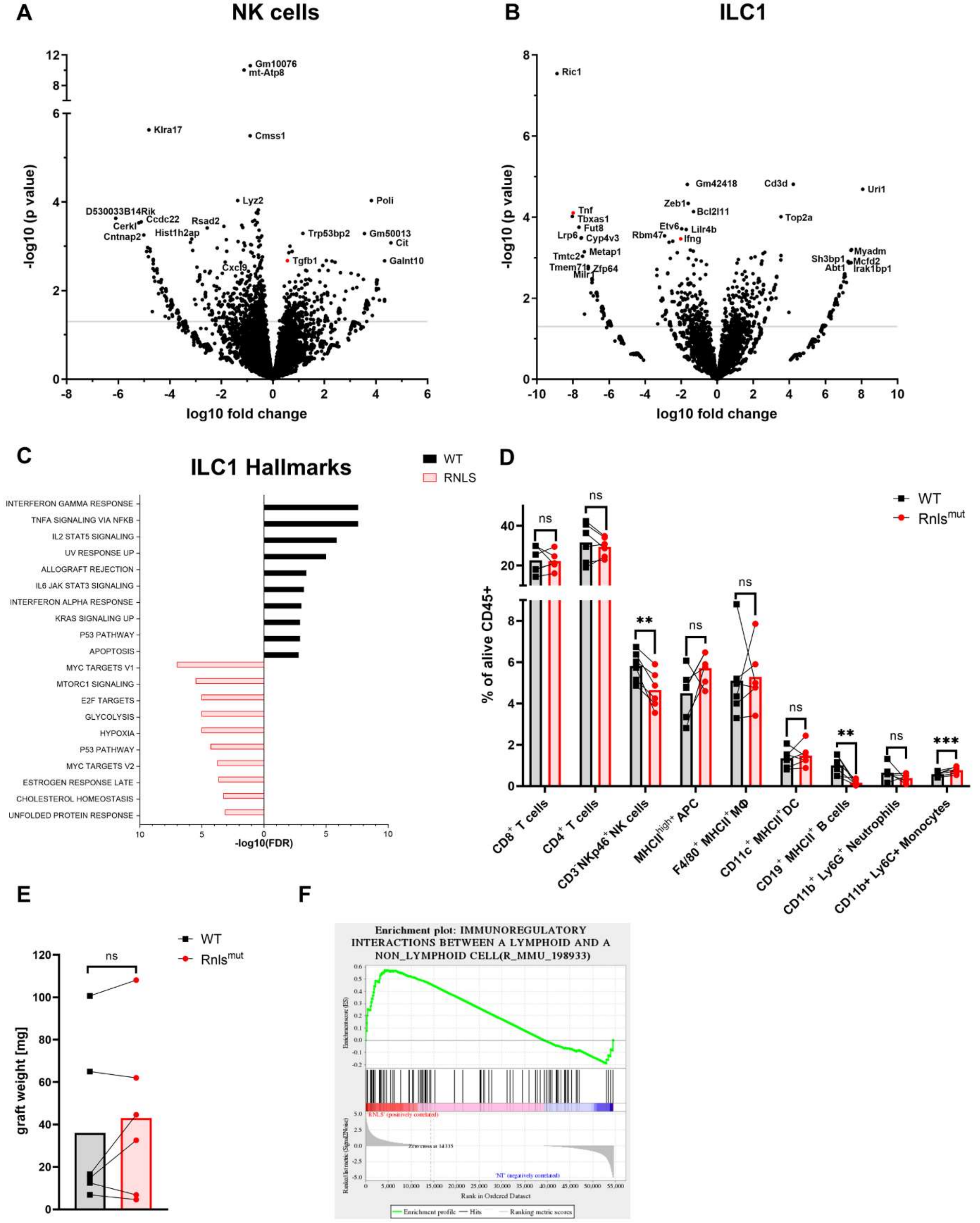

#### Supplemental Data Figure 2

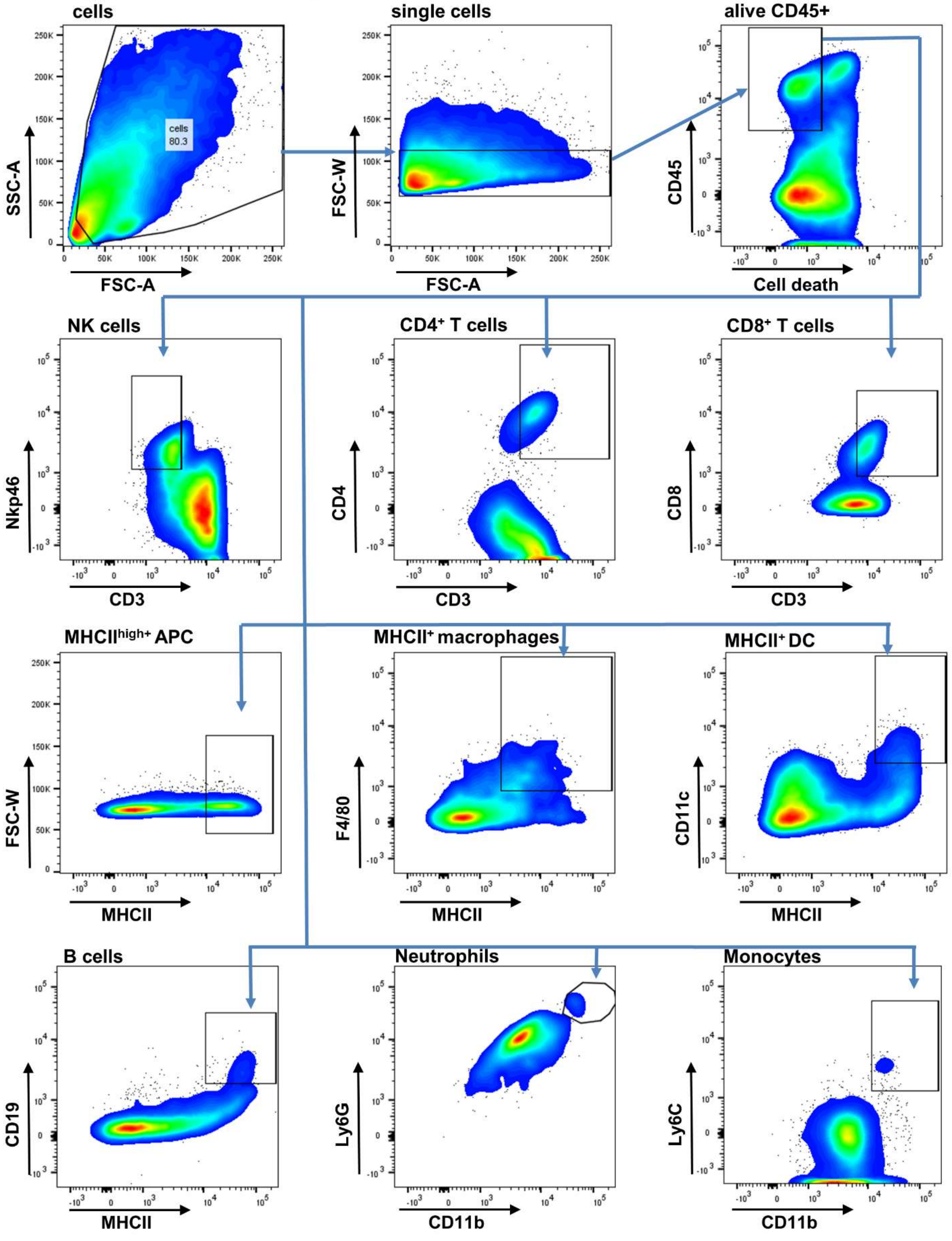

#### Supplemental Data Figure 3

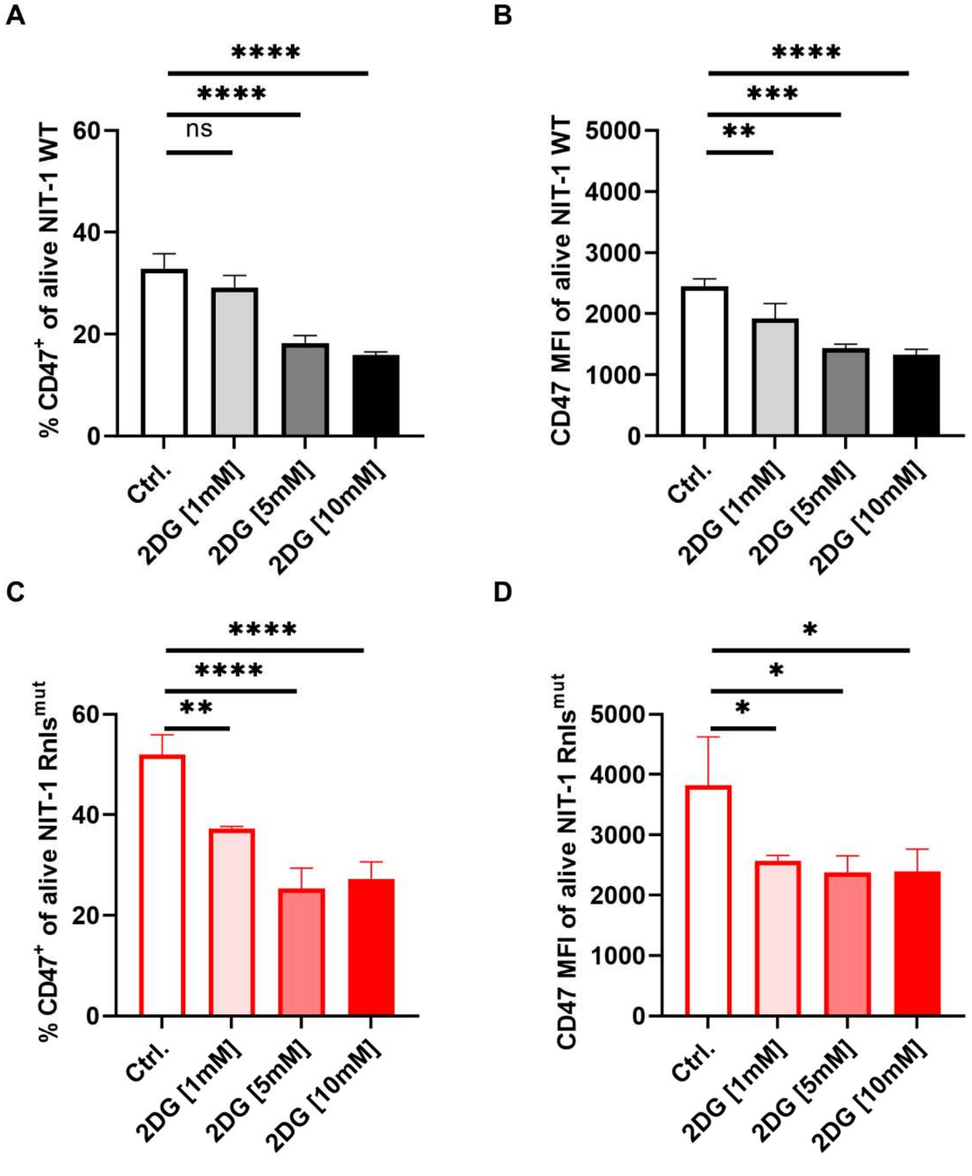

#### Supplemental Data Figure 4

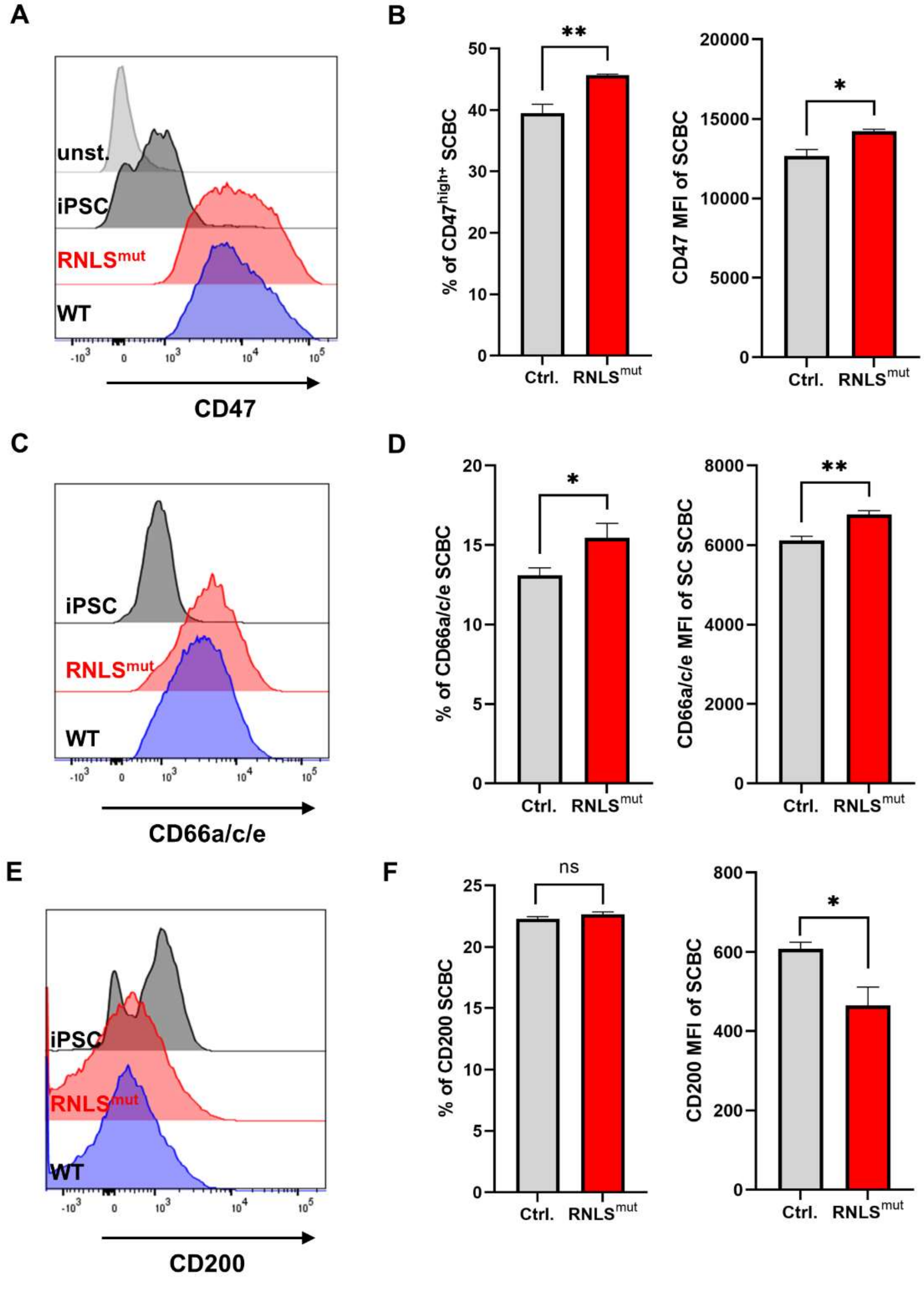
